# A cortical wiring space links cellular architecture, functional dynamics and hierarchies in humans

**DOI:** 10.1101/2020.01.08.899583

**Authors:** Casey Paquola, Jakob Seidlitz, Oualid Benkarim, Jessica Royer, Petr Klimes, Richard A. I. Bethlehem, Sara Larivière, Reinder Vos de Wael, Jeffery A. Hall, Birgit Frauscher, Jonathan Smallwood, Boris C. Bernhardt

## Abstract

The vast net of fibres within and underneath the cortex is optimised to support the convergence of different levels of brain organisation. Here we propose a novel coordinate system of the human cortex based on an advanced model of its connectivity. Our approach is inspired by seminal, but so far largely neglected models of cortico-cortical wiring established by post mortem anatomical studies and capitalizes on cutting-edge neuroimaging and machine learning. The new model expands the currently prevailing diffusion MRI tractography approach by incorporation of additional features of cortical microstructure and cortico-cortical proximity. Studying several datasets, we could show that our coordinate system robustly recapitulates established sensory-limbic and anterior-posterior dimensions of brain organisation. A series of validation experiments showed that the new wiring space reflects cortical microcircuit features (including pyramidal neuron depth and glial expression) and allowed for competitive simulations of functional connectivity and dynamics across a broad range contexts (based on resting-state fMRI, task-based fMRI, and human intracranial EEG coherence). Our results advance our understanding of how cell-specific neurobiological gradients produce a hierarchical cortical wiring scheme that is concordant with increasing functional sophistication of human brain organisation. Our evaluations demonstrate the cortical wiring space bridges across scales of neural organisation and can be easily translated to single individuals.

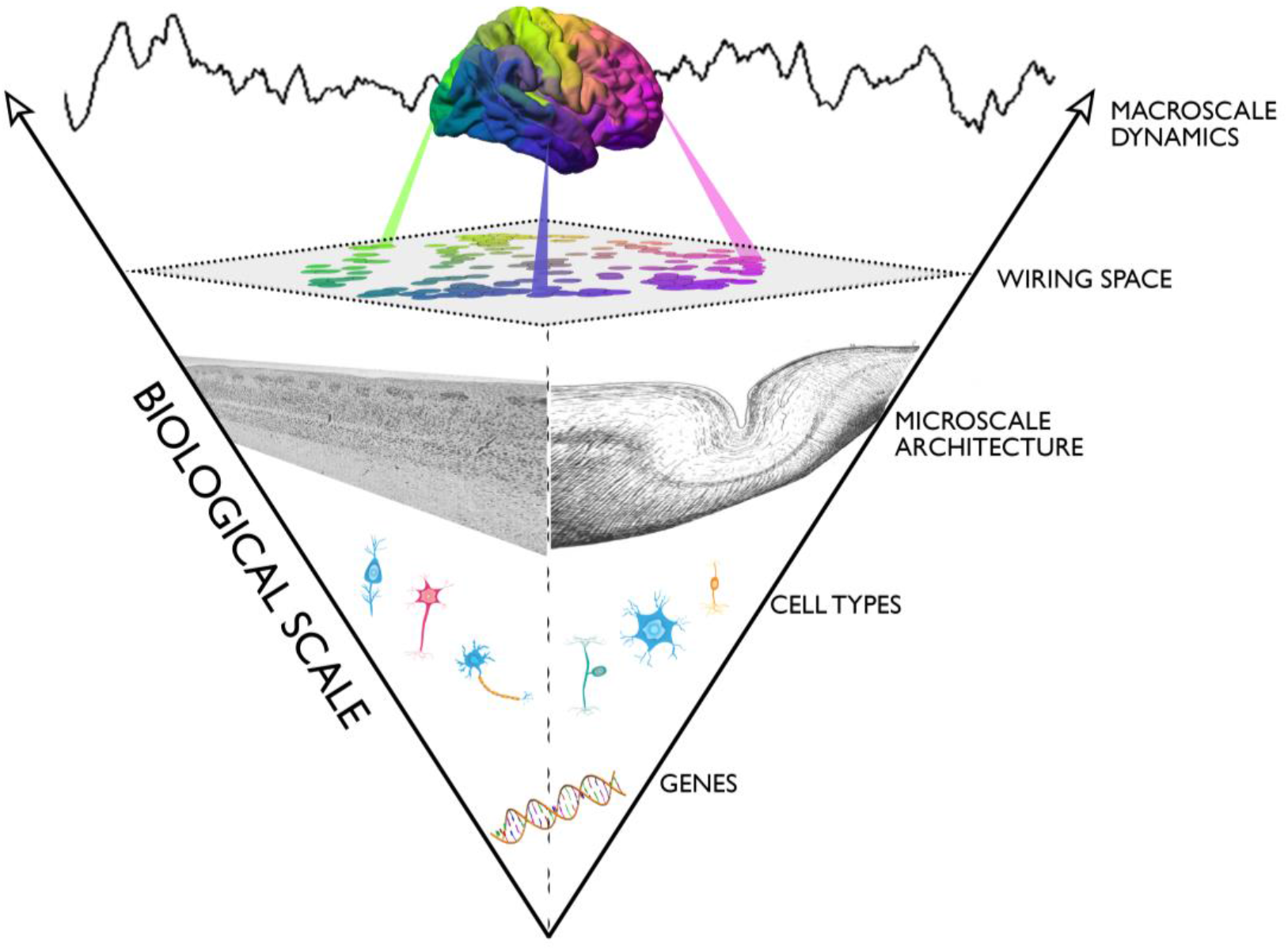

## Introduction

Neuronal activity in the cortex is simultaneously constrained by both local columnar circuitry and large-scale networks^1,2^. It is generally assumed that this interaction emerges because individual neurons are embedded in a global context through an intricate web of short- and long-range fibres. Developing a better model of this cortical wiring scheme is a key goal of systems neuroscience, because it would serve as a blueprint for the mechanisms through which local influences on neural function impact on spatially distant sites, and vice versa.

*Post mortem* histological and gene expression studies provide gold standard descriptions of how neurobiological and microstructural features are distributed across the cortex^3–6^. Histological and genetic properties of the brain often vary gradually together, mirroring certain processing hierarchies, such as the visual system^7^. This suggests that observed brain organisation may be the consequence of a set of consistent principles that are expressed across multiple scales (gene expression, cytoarchitecture, cortical wiring and macroscale function). Critically, full appreciation of how local and global features of brain organisation constrain neural function requires that multiple levels of brain organisation are mapped *in vivo.* To achieve this goal, our study capitalised on state-of-the-art magnetic resonance imaging (MRI) methods and machine learning techniques to build a novel model of the human cortical wiring scheme. We tested whether this model provides a meaningful description of how structure shapes macroscale brain function and the information flow between different systems. In particular, if our model successfully bridges the gap between micro- and macroscopic scales of neural organisation then it should describe local features of both cortical microcircuitry and its macroscale organisation and deliver meaningful predictions for brain function.

Currently, the prevailing technique to infer structural connectivity in living humans is diffusion MRI tractography^10,11^. By approximating white matter fibre tracts *in vivo*^10–12^, tractography has advanced our understanding of structural networks in health^13–15^, disease^16–18^, and shaped our understanding of the constraining role of brain structure on function^19–24^. Diffusion MRI tractography, however, has recognised limitations^25,26^. Crucially, the technique does not explicitly model intracortical axon collaterals and superficial white matter fibres, short-range fibres contributing to >90% of all cortico-cortical connections^27^. To address this gap, our approach combines diffusion tractography with two complementary facets of cortical wiring, namely spatial proximity and microstructural similarity. Spatial proximity predicts short cortico-cortical fibres^28,29^, which transmit the most common type of neural information also referred to as “nearest-neighbour-or-next-door-but-one”^30^. Microstructural similarity is a powerful predictor of interregional connectivity in non-human animals^31^, whereby the “structural model” of cortico-cortical connectivity postulates that connectivity likelihood between two regions is primarily governed by similarity in cytoarchitecture^32–35^. We recently developed and histologically validated microstructure profile covariance analysis, which quantifies microstructural similarity between different cortical areas *in vivo* through a systematic comparison of intracortical myelin sensitive neuroimaging profiles^7^. These complementary features can be fused using manifold learning techniques, resulting in a more holistic, multi-scale representation of cortical wiring. This extends upon previous work, in which we and others have derived manifolds from single modalities to map gradual changes in functional connectivity or tissue microstructure^7,36^.

Here, we generate a new coordinate system of the human cortex that is governed by complementary *in vivo* features of cortical wiring, expanding on traditional diffusion MRI tractography. Our wiring space incorporates advanced neuroimaging measures of cortical microstructure similarity, proximity, and white matter fibres, fused by non-linear dimensionality reduction techniques. We tested the neurobiological validity of our newly developed model by crossreferencing it against *post mortem* histology and RNA sequencing data^5,8,9^. Furthermore, we assessed the utility of our model to understand macroscale features of brain function and information flow by assessing how well the model predicts resting-state connectivity obtained from functional MRI as well as directed descriptions of neural function and processing hierarchy provided by intracerebral stereo-electroencephalography. These experiments were complemented with a comprehensive battery of robustness and replication analysis, to assess the consistency and generalizability of the new wiring model across analytical choices and datasets.

## Results

### A multi-scale model of cortical wiring

Cortical wiring was first derived from a *Discovery* subset of the Human Connectome Project dataset (HCP, n=100 unrelated adults) that offers high resolution structural magnetic resonance imaging (MRI), diffusion MRI, and microstructurally sensitive T1w/T2w maps^38^ (**Figure 1A**, see *Methods* for details).

**Figure 1.**
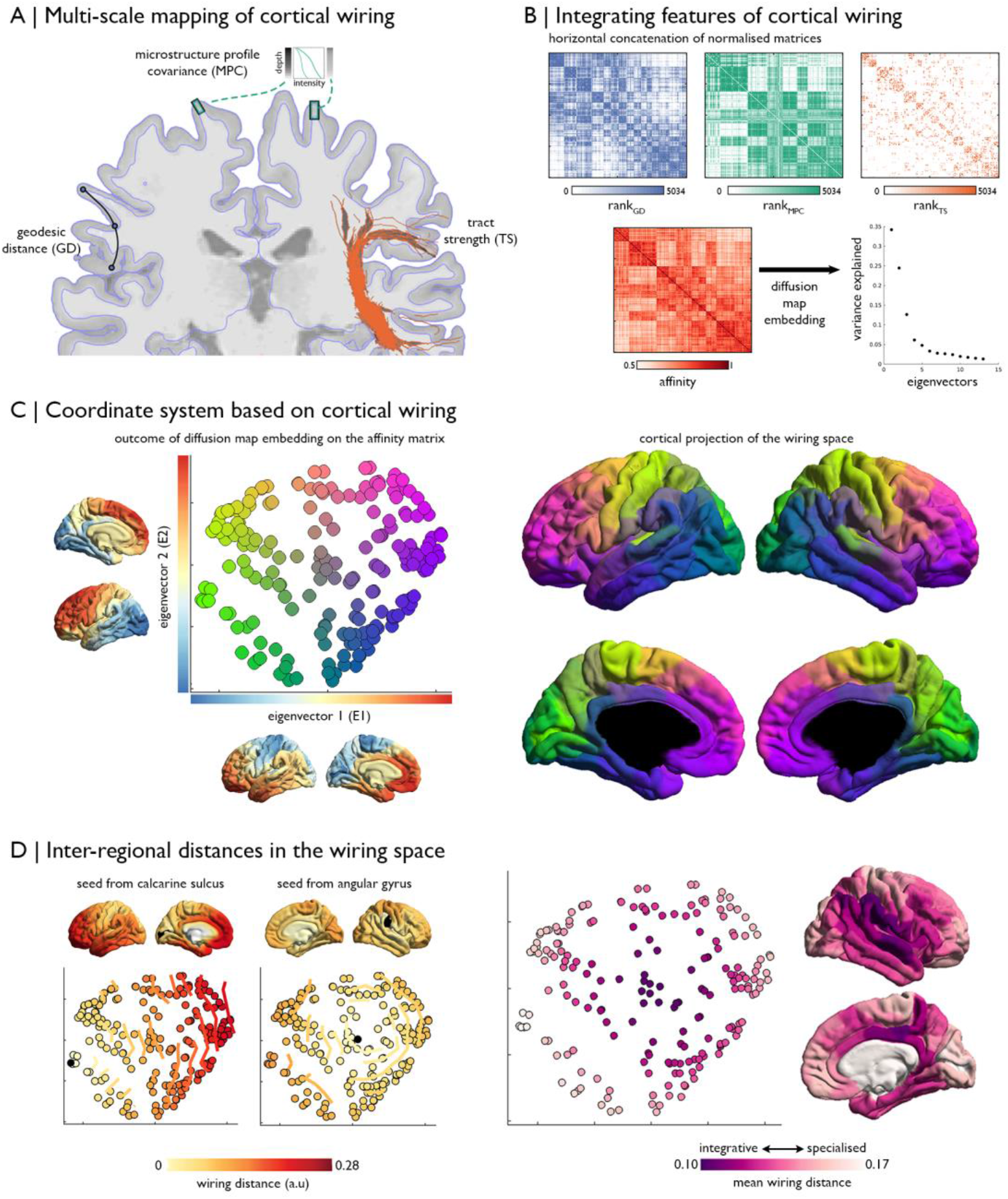
The multi-scale cortical wiring model. **(A)** Wiring features *i.e.,* geodesic distance (GD), microstructure profile covariance (MPC), and diffusion-based tractography strength (TS) were estimated between all pairs of nodes. **(B)** Normalised matrices were concatenated and transformed into an affinity matrix. Manifold learning identified a lower dimensional space determined by cortical wiring. **(c)** *Left.* Node positions in this newly-discovered space, coloured according to proximity to axis limits. Closeness to the maximum of the second eigenvector is redness, towards the minimum of the first eigenvector is greenness and towards the maximum of the first eigenvector is blueness. The first two eigenvectors are shown on the respective axes. *Right.* Equivalent cortical surface representation. **(D)** Calculation of inter-regional distances (isocontour lines) in the wiring space from specific seeds to other regions of cortex *(left).* Overall distance to all other nodes can also be quantified to index centrality of different regions, with more integrative areas having shorter distances to nodes *(right).* a.u.: arbitrary units.

The model integrated three complimentary features of structural connectivity, mapped between 200 spatially contiguous, evenly sized nodes: *i)* **Geodesic distance (GD)**, calculated as the shortest path between two nodes stepping through white or grey matter voxels, reflects the spatial proximity and cortico-cortical wiring cost of two regions^28^; *ii)* **Microstructure profile covariance (MPC)**, that is the correlation between myelin-sensitive imaging profiles taken at each node in the direction of cortical columns^7^, indexes architectonic similarity, the strongest predictor of projections in non-human primates^31^; *iii)* **Tract strength (TS)**, based on tractography applied to diffusion-weighted MRI, yields an estimate of the white matter tracts between each pair of nodes. Regional variations in the correspondence of wiring features, in terms of both magnitude and direction, highlight the necessity of multi-feature integration to provide a nuanced and expressive characterisation of a region’s structural connectivity (**Figure S1**).

To integrate these features into a compact coordinate system governed by cortical wiring, we normalised and concatenated the inter-regional matrices, computed an affinity matrix, and performed manifold learning (**Figure 1B**, see *Methods*). Diffusion map embedding, a nonlinear dimensionality reduction technique, was selected as a fast and robust approach that provides a global characterisation while preserving local structure in a data-driven manner^39^. Two dominant eigenvectors explained approximately 60% of variance in wiring affinity, with the first illustrating a sensory-fugal gradient (~35%) and the second an anterior-posterior gradient (~25%). These gradients represent principle axes of variation in cortical wiring (**Figure 1C**). The two-dimensional representation is hereafter referred to as the *wiring space*. Distances between two nodes in this new space provide a single integrative metric of cortical wiring affinity (**Figure 1D**). Nodes with high wiring affinity are close by, while dissimilar regions have a greater wiring distance. By taking the average wiring distance of each node, we found that the posterior cingulate, temporoparietal junction and superior temporal gyrus represent the integrative core of the wiring space (**Figure 1D**). In contrast, primary sensory areas, such as the calcarine sulcus and superior precentral gyrus, exhibit highly specialised cortical wiring.

To evaluate generalizability, we reconstructed the wiring model in an independent dataset of 40 healthy adults scanned at our imaging centre *(MICs* cohort; see *Methods* for details). While imaging parameters were comparable to the main cohort, this replication cohort involved acquisition of quantitative T1 relaxometry data to index intracortical microstructure instead of T1w/T2w maps^40–43^. Regardless of these site-wise idiosyncrasies, our procedure produced highly similar wiring spaces (correlations between both sites for eigenvectors 1/2: r=0.93/0.84, **Figure S2A**). The wiring space was also conserved at an individual level (**Figure S3A**). The most prominent inter-individual shifts in nodal positioning were observed in superior temporal and superior parietal regions (**Figure S3C**).

### Neurobiological underpinnings

We next evaluated the capacity of the new model to reflect local neurobiological features, by examining *post mortem* human histology and gene expression data. We generated cell-staining intensity profiles for each node from a high-resolution volumetric reconstruction of a single Merker stained human brain^8^ (**Figure 2A-C**) and extracted gene expression from mRNA sequencing data in eleven neocortical areas, each matched to one node^9,44^ (**Figure 2D-F**). Cytoarchitectural similarity and gene co-expression were correlated to wiring distance (histology: r=−0.53, p<0.001 **Figure 2B**; co-expression: r=-0.64, p<0.001). Furthermore, the above associations were robust even when additionally correcting for the influence of physical distance along the cortex (histology: partial r=-0.48, p<0.001; co-expression: partial r=-0.38, p<0.001).

**Figure 2.**
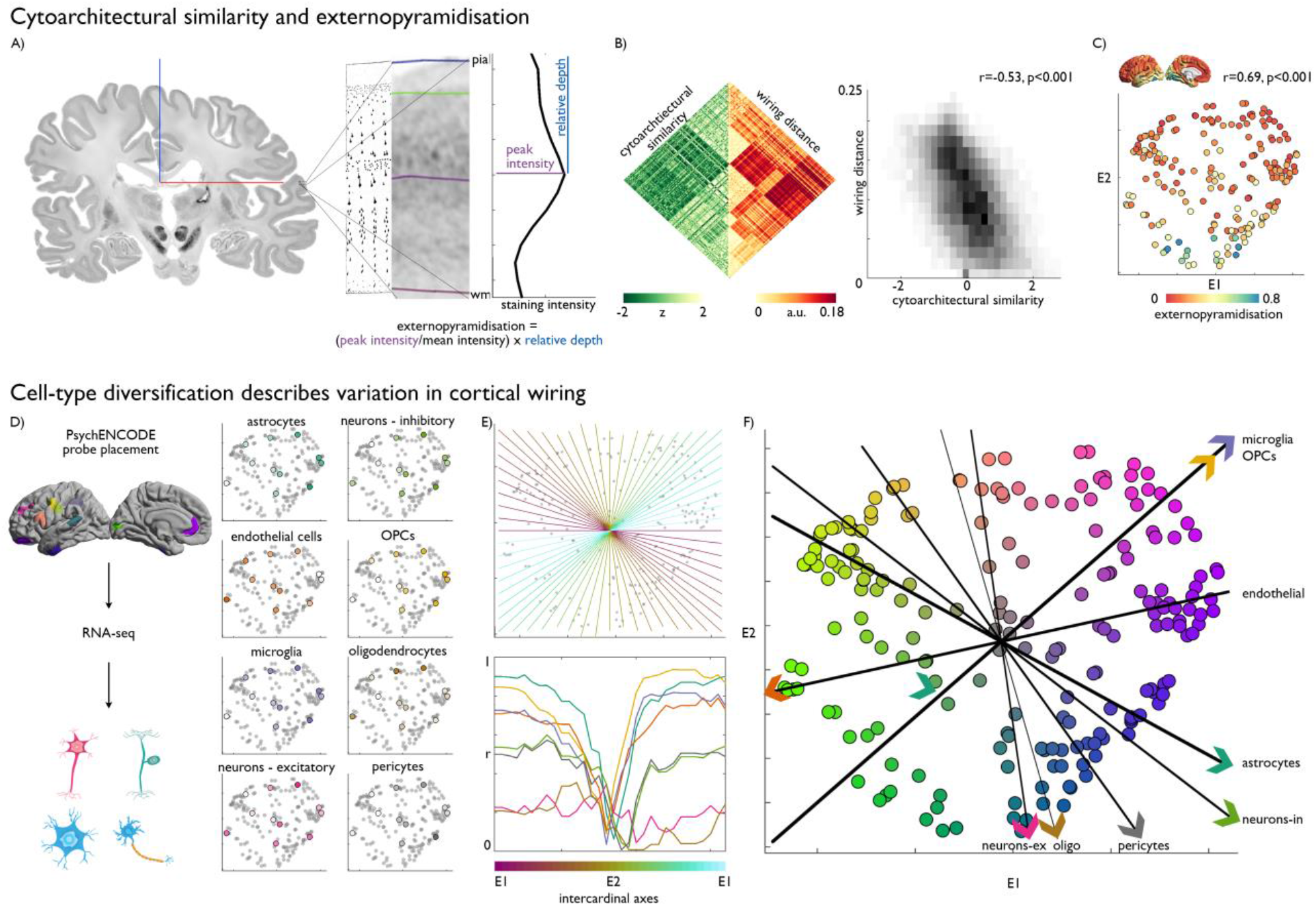
Neurobiological substrates of the wiring space. **(A)** A 3D *post mortem* histological reconstruction of a human brain^8^ was used to estimate cytoarchitectural similarity and externopyramidisation. Here, we present a coronal slice, a drawing of cytoarchitecture^46^, magnified view of cortical layers in BigBrain and a staining intensity profile with example of calculation of externopyramidisation^45^. **(B)** Matrix and density plot depict the correlation between BigBrain-derived cytoarchitectural similarity and wiring distance between pairs of regions. **(c)** Externopyramidisation projected onto the cortical surface and into the wiring space. **(D)** mRNA-seq probes, assigned to eleven representative nodes (coloured as in **Figure 1C**, i.e. their position in the wiring space), provided good coverage of the space and enabled characterisation of cell-type specific gene expression patterns. Average cell-type specific gene expression patterns projected in the wiring space, with brighter colours signifying higher expression. **(E)** Equally spaced intercardinal axes superimposed on the wiring space, and below, line plots showing correlation of gene expression patterns with each of the axes. **(F)** Strongest axis of variation *(i.e.,* maximum |r|) for each cell-type.

We hypothesised that the principle axes of the cortical wiring scheme would describe systematic variations in cytoarchitecture that reflect a region’s position in a neural hierarchy. It has previously been proposed that externopyramidisation is optimally suited to assess hierarchy-dependent cytoarchitecture because it tracks the laminar origin of projections, which signifies the predominance of feedback or feedforward processing^5^. Externopyramidisation was estimated from histological markers capturing the relative density and depth of pyramidal neurons^45^ (**Figure 2A**), the primary source of interregional projections. Increasing values reflect a shift from more infragranular feedback connections to more supragranular feedforward connections^5,45^. Multiple linear regression indicated that the eigenvectors of the wiring space explained substantial variance of externopyramidisation (r=0.69, pspin<0.001; **Figure 2C**) and was independent of regional variations in cortical morphology (**Figure S4A**). Notably, the complete model that aggregated all wiring features *(i.e.,* MPC, GD, TS) in a low dimensional space explained more variance in externopyramidisation than models constructed using alternative combinations or subsets of the wiring features (**Table S1**). Externopyramidisation gradually decreased along the second eigenvector, suggesting that our wiring space captures a posterior to anterior transition from feedforward to feedback processing.

We further explored how the broader cellular composition and microcircuitry relates to the layout of the wiring space. To do so, we estimated the expression of eight canonical cell-type gene sets in eleven cortical areas (see *Methods)* and found that these expression patterns accounted for significant variance in the macroscale organisation of cortical wiring (up to R^2^=0.83, **Figure 2D**, **Table S2**). By defining the strongest axis of variation for each cell-type in the wiring space, we discovered distinct spatial gradients of the cell-types, which together depicted the multiform cellular differentiation of the wiring space (**Figure 2E-F**). These findings established that the wiring space captures the organisation of neuronal and non-neuronal cells (**Figure 2F**), and offers a new line of evidence on the heightened expression of neuro-modulatory glia, such as astrocytes and microglia^47,48^, towards the transmodal areas.

### Constraining role for functional architecture

Thus far our analysis indicates that the wiring space successfully captures macroscale spatial trends in cortical organisation, and that is reflects underlying cytoarchitectonic and cellular microcircuit properties. Next, we tested the hypothesis that our wiring space also underpins the functional architecture of the brain using a series of analyses.

First, we mapped well established intrinsic functional communities^49^ into the wiring space and inspected their relative distances (**Figure 3A-C**). We found that sensory networks were located in the lower (visual) and upper (somatomotor) left extremities of the new coordinate system. Sensory nodes spanned a broad range of wiring distances, reflecting transitions from specialised to integrative connectivity. In contrast, transmodal default, frontoparietal and limbic networks were located more towards the right extremities in the wiring space. The dorsal attention network created a frontier between sensory networks and the transmodal extremity, while the ventral attention also subsumed an intermediary position at the integrative core. Together, these analyses demonstrate how the relative positioning of functional communities in the structural wiring space underpins coordination within and between functional networks.

**Figure 3.**
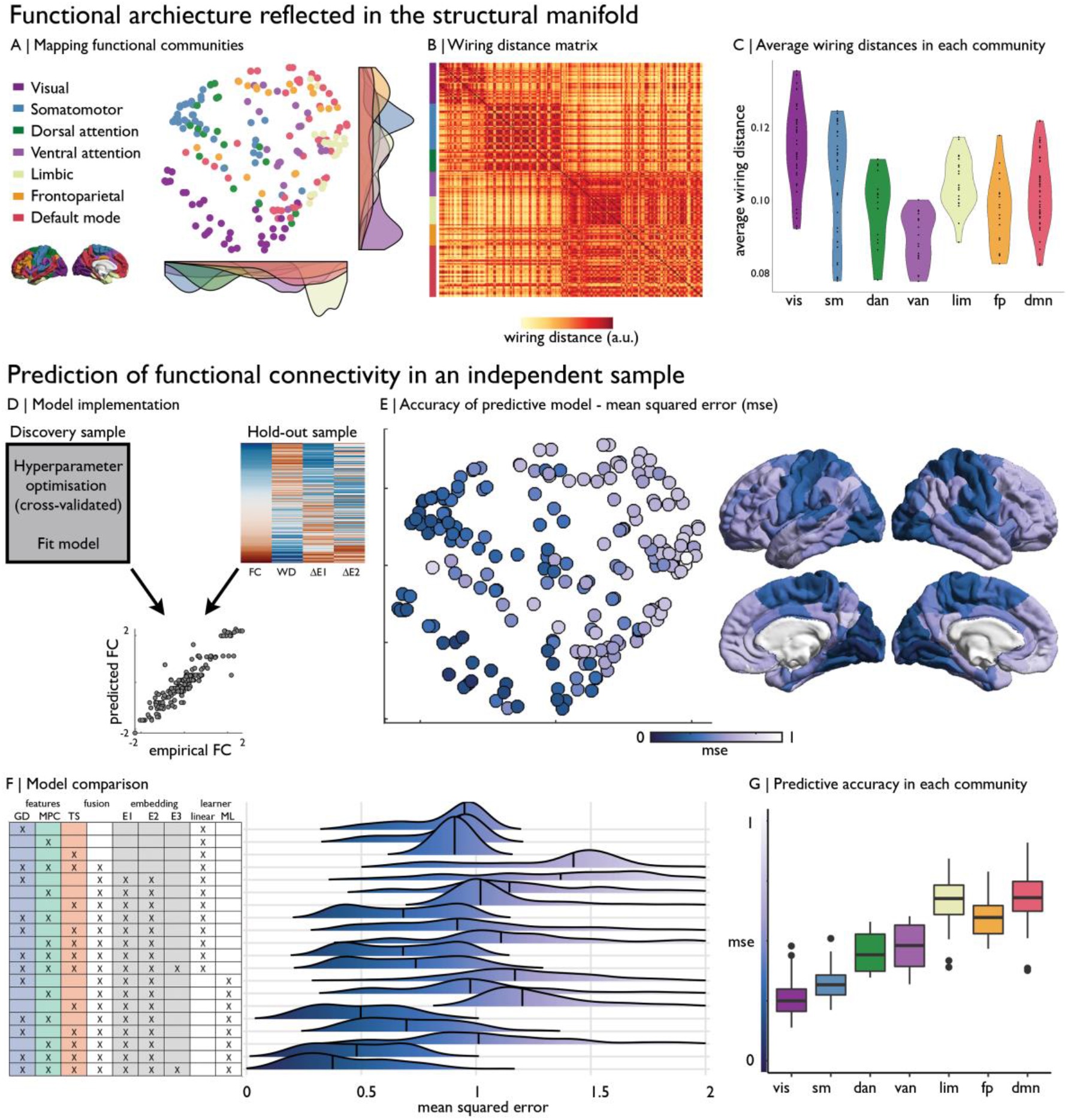
From cortical wiring to functional connectivity. **(A)** Nodes in the wiring derived coordinate system coloured by functional community^49^, with the distribution of networks shown by density plots along the axes. **(B)** Wiring distance between nodes, ordered by functional community, revealed a modular architecture. **(C)** Violin plots show the average wiring distance for nodes in each functional community, with higher values being more specialised in their cortical wiring. **(D)** Using the boosting regression models from the *Discovery* dataset, we used features of the wiring space to predict z-standardised functional connectivity in a *Hold-out* sample. The model was enacted for each node separately. FC: functional connectivity. WD: wiring distance. ΔE1: difference on eigenvector 1. ΔE2: difference on eigenvector 2. **(E)** Mean squared error across nodes are shown in the wiring space and on the cortical surface. **(F)** Predictive accuracy of various cortical wiring models, involving the use of different features, multi-feature fusion, eigenvectors from diffusion map embedding and a linear or machine learning (ML) learner. **(G)** Mean squared error of the wiring space model stratified by functional community.

Secondly, we assessed whether our wiring space can also predict macroscale functional brain connectivity. We used a supervised machine learning paradigm that applies adaptive boosting to predict functional connectivity based on relative distances of nodes in the wiring space (**Figure 3D;** see *Methods).* The new space, trained on the *Discovery* dataset, predicted resting-state functional connectivity in the independent *Hold-out* sample with high accuracy (mean squared error=0.49±0.19; R^2^=0.51±0.20; **Figure 3E**), and outperformed learners trained on data from fewer cortical wiring features or learners trained on all modalities but without using manifold embedding (**Figure 3F**). Inspecting regional variations in predictive accuracy indicated that cortical wiring topography was more tightly linked to functional connectivity in sensory areas; systems upon which classical examples of the cortical hierarchy were developed^50^, while it tapered off towards transmodal cortex. Further expanding the wiring space to three eigenvectors/dimensions mildly decreased the average mean squared error, but resulted in larger variance across nodes and poorer predictive accuracy at the lower limits for the prediction of resting-state functional connectivity (mean squared error=0.44±0.21; R^2^=0.56±0.22). While accuracy was reduced, the two-dimensional model also provided state-of-the-art predictions of resting-state functional connectivity in individual participants of the *Hold-out* dataset (mean squared error=0.89±0.24, R^2^=0.16±0.17; **Figure S3B**).

High predictive performance at the group- and individual subject level could be replicated in the independent *MICs* dataset, despite the smaller sample size (group-level split half testing mean squared error= 0.63±0.22, R^2^=0.37±0.22; **Figure S2B;** individual level: mean squared error= 1.06±0.46, R^2^=0.11±0.14).

One important aspect of a structural wiring scheme may be its potential ability to simulate a broad functional range. As an additional proof of concept for the wiring model to be such a versatile framework, we showed that the wiring model could also predict functional connectivity across different task conditions, in both HCP and MICs subsamples *(HCP:* 7 tasks: mean squared error=0.64±0.22, R^2^=0.36±0.22 **Figure S5**. *MICs:* 3 tasks: mean squared error=0.89±0.21, R^2^=0.16±0.21).

### Large-scale organisation of directed coherence

The above analyses showed that the wiring space robustly explains substantial aspects of macroscale functional organisation and connectivity. We next examined whether it can also account for a more direct measure of neural functional connectivity, by examining stereo-electroencephalographic recordings during resting wakeful rest in ten epileptic patients (who underwent multimodal imaging before the implantations, with imaging identical to the *MICs* dataset; **Figure 4A**). Patients presented with a similar wiring space solution as controls from the same sample (**Figure S6;** correlations between eigenvectors 1/2: r=0.83/0.82). In line with the above functional connectivity analysis, the wiring model explained substantial within sample variance in undirected coherence (R^2^=0.60±0.23; **Figure 4B**; **Figure S7**), especially in frequencies >15Hz 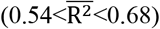. We calculated the phase slope index within frequency bands as an estimate of unidirectional flow^51^. To highlight large-scale organisation and account for the incomplete coverage of electrodes in each participant, we clustered the wiring space into twelve macroscopic compartments (**Figure 4C**; see *Methods* for determination of k=12). We estimated the phase slope index between each pair of clusters and performed significance testing using a linear mixed effect model that included subject as a random effect. Decomposition of inter-cluster similarities in the phase slope index using a principle component analysis revealed a gradient running across the wiring space. The first principle component, accounting for 39% of variance, illustrated a transition in the patterns of directed coherence from the upper left to lower right of the wiring space, that is running from central to temporal and limbic areas (**Figure 4D**). The component loading was underpinned by varied expression of cell-types (**Figure 4D**). Lasso regularisation showed that inhibitory neuron expression was the most important cellular feature in supporting this coherence-derived topography, followed by endothelial cells in highly regularised models. Together, inhibitory neuron and endothelial cell expression accounted for 44% of variance in component loading (p=0.04). The cell-types did not reach significance in explaining the component loading independently, emphasising the multivariate contribution of cell-types to spatial variations in electrophysiological oscillations. Robustness of the component loading and edge-wise phase slope index estimates were supported by a leave-one-subject-out procedure (**Figure S7**).

**Figure 4.**
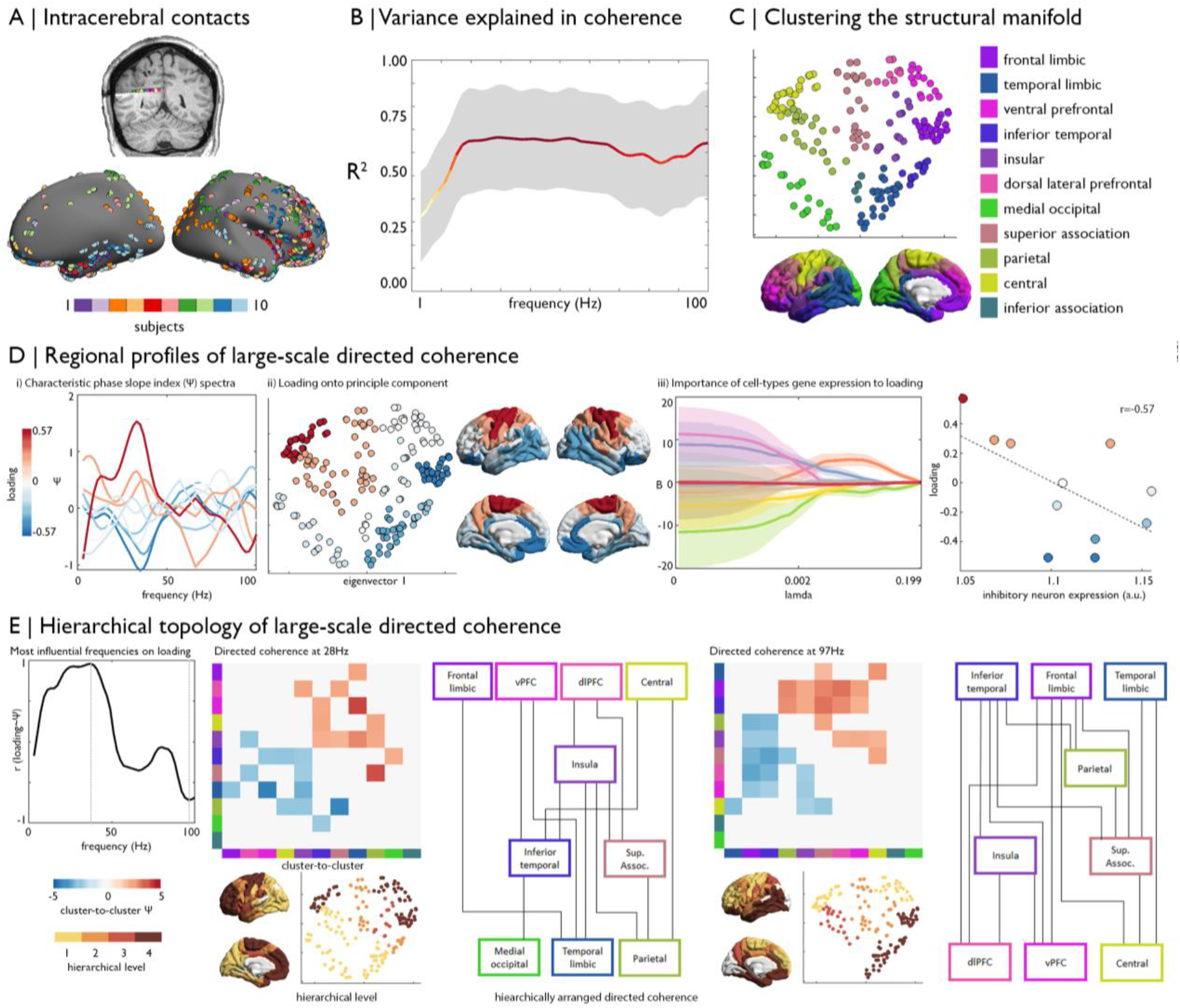
Hierarchical information processing is organised along the wiring space. **(A)** Intracerebral implantations of ten epileptic patients were mapped to the cortical surface, and intracortical EEG contacts selected. We studied five minutes of wakeful rest. **(B)** Mean and standard deviation in the variance explained in undirected coherence by wiring space features using adaboost machine learning across all nodes. **(C)** Clusters of the wiring space. **(D)** Phase slope index (Ψ) was calculated for each pair of intra-subject electrodes, then cluster-to-cluster estimates were derived from a linear mixed effect model. Pearson correlation across Ψ estimates was used to measure the similarity of clusters, and the major axis of regional variation was identified via principle component analysis. *i)* Average Ψ spectra for each region coloured by loading on the first principle component (accounting for 39% of variance). *ii)* Component loadings presented in the wiring space and on the cortical surface illustrate a gradient from upper left regions, corresponding to central areas, toward lower right areas, corresponding to temporal and limbic areas. *iii)* Lasso regularisation demonstrates the contribution of cell-type specific gene expression (colours matching **Figure 2**) and externopyramidisation (red) to explain variance in the component loadings. For example, inhibitory neurons expression levels (green) are closely related to the component loading, as shown in the scatterplot. Shaded areas show the standard deviation in fitted least-squares regression coefficients across leave-one-observation-out iterations. **(E)** The most influential frequencies on the component loading were identified through a correlation of average Ψ spectra (**Figure 4Di**) with the component loading. For the global maxima and minima, we present cluster-to-cluster Ψ estimates, thresholded at p<0.05, suprathreshold edges plotted as a hierarchical schema and the hierarchical level of each cluster on the cortical surface and in the wiring space.

Finally, we examined the topology of large-scale networks of directed coherence in frequencies influencing the component loading, and specifically tested whether they met the criteria for hierarchical organisation ^50^ (**Figure 4E**). Both the 26-30Hz (beta) and 95-99Hz (high gamma) bands met criteria for hierarchy, insomuch that clusters could be placed in levels that depict unidirectional flow of oscillations from the top to bottom of the graph. The beta band was associated with an anterior-posterior wave, whereas high gamma band was related to limbic-to-prefrontal oscillations. These results support the hierarchical organisation of large-scale directed coherence, which propagate as waves of oscillations moving through the cortical wiring scheme, and also demonstrate the co-occurrence of hierarchies and that these are operationalised in different frequencies.

## Discussion

Based on advanced machine learning of multiple features sensitive to cortico-cortical wiring, our work identified a novel and compact coordinate system of human cortex. Our analysis established that cortical wiring is dominated by two principal axes, one running from sensory towards transmodal systems and one running from anterior to posterior. Critically, this novel space successfully accounted for both local descriptions of cortical microcircuitry as well as macroscale cortical functional dynamics measured by functional MRI and intracranial electrical recordings. By projecting *post mortem* histological and transcriptomic profiles into this newly discovered space, we could demonstrate how these axes are determined by intersecting cell-type specific and cytoarchitectural gradients. In addition to establishing these local neurobiological features, our findings support that the wiring-based space serves as a powerful scaffold within which macroscale cortical function can be understood. Using both non-invasive imaging in healthy individuals, and direct neuronal measurements in a clinical population, we demonstrated that the wiring-derived manifold describes how neural function is hierarchically organised in both space and time. Our findings were replicable in different datasets and at the single subject level; moreover, a series of additional experiments showed that this novel representation outperformed conventional approximations of structural connectivity in their ability to predict function. Together, we have successfully identified a compact description of the wiring of the cortex that can help to ultimately understand how neural function is simultaneously constrained by both local and global features of cortical organisation.

Our multivariate model of cortical wiring reflects an important extension on diffusion MRI tractography because it additionally incorporates spatial proximity and microstructural similarity, both features tap into generative principles of cortico-cortical connectivity as demonstrated by prior human and non-human animal studies^32–35^. The application of manifold learning to this enriched representation of cortical wiring helped to determine a low dimensional, yet highly expressive, depiction of cortical wiring. In other fields, notably genomics and data science more generally, embedding techniques have become widely adopted to identify and represent the structure in complex, highdimensional datasets^9,39,52,53^. In recent neuroimaging studies, several approaches have harnessed non-linear dimensionality reduction techniques to identify manifolds from single modalities, highlighting changes in microstructure and function at the neural system level^7,36,54,55^. The wiring space identified here captured both sensory-fugal and anterior-posterior processing streams, two core modes of cortical organisation and hierarchies established by seminal tract-tracing work in non-human primates^37,56^. The anterior-posterior axis combines multiple local gradients and functional topographies, such as the ventral visual stream running from the occipital pole to the anterior temporal pole that implements a sensory-semantic dimension of perceptual processing^57,58^ and a rostro-caudal gradient in the prefrontal cortex that describes a transition from high-level cognitive processes supporting action preparation to those tightly coupled with motor execution^56,59–61^. The sensory-fugal axis represents an overarching organisational principle that unites these local processing streams. While consistent across species, the number of synaptic steps from sensory to higher-order systems has increased throughout evolution, supporting greater behavioural flexibility^37^ and decoupling of cognitive functioning from the here and now^58^. By systematically studying inter-nodal distances within the wiring derived space, it may be possible to gain a more complete understanding of the difference between specialised systems at the periphery and more centrally localised zones of multimodal integration, such as the temporoparietal junction and cingulate cortex. Many of these regions have undergone recent evolutionary expansion^62^, are sites of increased macaque-human genetic mutation rates^63^, and exhibit the lowest macaque-human functional homology^64^. In this context, the more complete model of cortical wiring provided here may play a critical role in advancing our understanding of how changes in cortical organisation have given rise to some of the most sophisticated features of human cognition.

Our new coordinate system reflects an intermediate description of cortical organisation that simultaneously tracks its microstructural underpinnings and also addresses the emergence of functional dynamics and hierarchies at the system level. Cross-referencing the new space with a 3D histological reconstruction of a human brain^8^ established close correspondence between our *in vivo* model and histological measurements. This work adds to the notion that cytoarchitecture and cortical wiring are inherently linked^65,66^, as variations in projections across cortical layers determine the layout of the cortical microcircuitry^67^. Feedforward connections often originate in supragranular layers (and terminate in lower layers in the target regions), while infragranular layers give rise to feedback projections flowing down the cortical hierarchy^68–71^. In this work, we observed an alignment between the cortical wiring scheme and a proxy for externopyramidisation^45^, which was sensitive to inter-areal differences in the depth of peak pyramidal neuron density that co-occurs with shifts from feedback to feedforward dominated connectivity profiles^5^. In addition to showing cytoarchitectonic underpinnings of the cortical microcircuitry, we capitalised on *post mortem* gene expression datasets derived from mRNA sequencing^72^, a technique thought to be more sensitive and specific than microarray transcriptomic analysis. This approach revealed that divergent gradients of cell-type specific gene expression underpin intercardinal axes of the new coordinate system, particularly of non-neuronal cell-types. Increased glia-to-neuron ratios in transmodal compartments of the new space may support higher-order cognitive functions, given comparative evidence showing steep increases in this ratio from worms to rodents to humans^73–76^. Astrocytes, in particular, exhibit morphological variability that may lend a cellular scaffold to functional complexity and transmodal processing^73^. For example, a uniquely human inter-laminar astrocyte was recently discovered with long fibre extensions, likely supporting long-range communication between distributed areas that may contribute to flexible, higher-order cognitive processing^77^.

Our coordinate system also established how structural constraints relate to cortical dynamics and information flow throughout hierarchical and modular systems. We showed that functional communities are circumscribed within the wiring derived space, supporting dense within-network connectivity, and that their relative positions describe a progressive transition from specialised sensory wiring to an integrative attentional core and distributed transmodal networks. Region-to-region distances in the wiring space provided competitive predictions of resting-state functional connectivity data, both at the level of the group and of a single subject. Spatial proximity and microstructural similarity are critical elements of the predictive value of our models, highlighting intracortical and cytoarchitecturally-matched projections in shaping intrinsic functional organization. However, these dominant aspects of cortical wiring are often not considered by computational models that simulate functional connectivity from measures of diffusion-based tractography^19,20,78^. In addition to feature enrichment, the use of nonlinear dimensionality reduction enhanced the predictive performance by minimising the influence of noisy edges and magnifying effects from the most dominant axes of cortical wiring. Such a model led to maximal gains in predictive power in unimodal areas, likely owing to their more locally clustered, and hierarchically governed connectivity profiles^79–81^. Predictive performance decreased towards transmodal networks, a finding indicating that more higher-level systems may escape (currently measurable) structural constraints and convergence of multisynaptic pathways^82^. Such conclusions are in line with recent work showing that transmodal areas exhibit lower microstructure-function correspondence^7^ and reduced correlations between diffusion tractography and resting state connectivity^83,84^, potentially contributing to greater behavioural flexibility^7,85^. The hierarchical nature of the wiring space was further supported by analysing its correspondence to direct measurements of neural dynamics and information flow via intracerebral stereo-electroencephalography. The sources and relative distances of beta and gamma hierarchies align with previous studies. Beta oscillations emanate from the sensorimotor cortex, supporting the maintenance of the current sensorimotor set^86,87^, and time lagged coherence during rest occurs between distal areas^88,89^. In contrast, gamma oscillations are robustly associated with a limbic source and, nested in theta oscillations, facilitate limbic-prefrontal coupling^88,90–92^. Using a strict definition of hierarchy as the topological sequence of projections^50,93^, we demonstrated that the wiring space underpins large-scale, frequency-specific waves of oscillations propagating from anterior to posterior and from limbic to prefrontal. Furthermore, bridging across scales, we provide an additional line of evidence for the importance of inhibitory neurons for supporting specific frequencies of oscillations, such as the role of somatostatin for beta oscillations^94^. One key feature of the present framework, therefore, is that it provides a basis to quantitatively assess how the interplay of neuronal oscillations underpin complex cortical organisation. Even with limited spatial resolution, data-driven decomposition of the wiring space proved an effective model of electrophysiological organization, in line with recent work showing the modular architecture of electrocorticography^95^.

Future studies should increase the resolution of the wiring space, which may be possible with ongoing efforts to generate robust estimates of white matter tracts from single voxels^96^. In lieu of a gold standard for cortical wiring in humans, the present work focused on equally balanced cortical wiring features, however, supervised learning techniques could reveal their relative importance for specific tasks or scales. As our results have shown, this novel representation of cortical wiring provides a practical workspace to interrogate the coupling of brain structure and function, and to study links between microcircuit properties and macroscale hierarchies. As such, the wiring space can be a powerful tool for to study the mutli-scale complexities of brain development, aging and disease.

## Methods

### Human Connectome project dataset

#### a) Data Acquisition

We studied data from 197 unrelated healthy adults from the S900 release of the Human Connectome Project (HCP; Glasser et al., 2013). The *Discovery* dataset included 100 individuals (66 females, mean±SD age=28.8±3.8 years) and the *Hold-out* dataset included 97 individuals (62 females, mean±SD age=28.5±3.7 years). MRI data were acquired on the HCP’s custom 3T Siemens Skyra equipped with a 32-channel head coil. Two T1w images with identical parameters were acquired using a 3D-MPRAGE sequence (0.7mm isotropic voxels, matrix=320×320, 256 sagittal slices; TR=2400ms, TE=2.14ms, TI=1000ms, flip angle=8°; iPAT=2). Two T2w images were acquired using a 3D T2-SPACE sequence with identical geometry (TR=3200ms, TE=565ms, variable flip angle, iPAT=2). A spin-echo EPI sequence was used to obtain diffusion weighted images, consisting of three shells with *b*-values 1000, 2000, and 3000s/mm^2^ and up to 90 diffusion weighting directions per shell (TR=5520ms, TE=89.5ms, flip angle=78°, refocusing flip angle=160°, FOV=210×180, matrix=178×144, slice thickness=1.25mm, mb factor=3, echo spacing=0.78ms). Four rs-fMRI scans were acquired using multi-band accelerated 2D-BOLD echo-planar imaging (2mm isotropic voxels, matrix=104×90, 72 sagittal slices, TR=720ms, TE=33ms, flip angle=52°, mb factor=8, 1200 volumes/scan). Participants were instructed to keep their eyes open, look at fixation cross, and not fall asleep. Seven task-evoked fMRI scans (working memory, gambling, motor, language, social cognition, relational processing and emotion processing^97^) were acquired with the same echo-planar imaging sequence as rs-fMRI with a total run time of 24:23 (min:sec). While T1w, T2w, diffusion scans and task-fMRI were acquired on the same day, rs-fMRI scans were split over two days (two scans/day).

#### b) Data preprocessing

MRI data underwent HCP’s minimal preprocessing^98^. Cortical surface models were constructed using Freesurfer 5.3-HCP^99–101^, with minor modifications to incorporate both T1w and T2w^102^. Following intensity nonuniformity correction, T1w images were divided by aligned T2w images to produce a single volumetric T1w/T2w image per subject, a contrast ratio sensitive to cortical microstructure (Glasser and Van Essen, 2011). Diffusion MRI data underwent correction for geometric distortions and head motion^98^. BOLD timeseries were corrected for gradient nonlinearity, head motion, bias field and scanner drifts, then subjected to ICA-FIX for removal of additional noise^103^. The rs-fMRI data were transformed to native space and timeseries were sampled at each vertex of the MSMAll registered midthickness cortical surface^104,105^.

#### c) Generation of the wiring features

Cortical wiring features were mapped between 200 spatially contiguous cortical ‘nodes’. The parcellation scheme preserves the boundaries of the Desikan Killany atlas^106^ and was transformed from fsaverage7 to subject-specific cortical surfaces via nearest neighbour interpolation.

##### Geodesic distance

Geodesic distance (GD) was calculated across subject-specific mid-cortical surface maps in native space. Exemplar vertices of each node were defined for each subject as the vertex with minimum Euclidian distance to the subject-specific node centroid. For each node, we matched the exemplar vertex to the nearest voxel in volumetric space, then used a Chamfer propagation (imGeodesics Toolbox; https://github.com/mattools/matImage/wiki/imGeodesics) to calculate the distance to all other voxels travelling through a grey/white matter mask. This approach differs from previous implementations of geodesic distance^28,29,36^ by involving paths through the grey and white matter, allowing for jumps within gyri and interhemispheric projections^107^. We projected GD estimations back from volumetric to surface space, averaged within node and produced a 200×200 GD matrix.

##### Microstructure profile covariance

The full procedure of the MPC approach may be found elsewhere^7^. In brief, we generated 12 equivolumetric surfaces between the outer and inner cortical surfaces^108^, and systematically sampled T1w/T2w values along linked vertices across the whole cortex. T1w/T2w intensity profiles were averaged within nodes, excluding outlier vertices with median intensities more than three scaled median absolute deviations away from the node median intensity. Nodal intensity profiles underwent pairwise Pearson product-moment correlations, controlling for the average whole-cortex intensity profile. The MPC matrix was absolutely thresholded at zero; remaining MPC values were then log-transformed to produce a symmetric 200×200 MPC matrix.

##### Tract strength

Tractographic analysis was based on MRtrix3 (https://www.mrtrix.org). Response functions for each tissue type were estimated using the dhollander algorithm^109^. Fibre orientation distributions were modelled from the diffusion-weighted MRI with multi-shell multi-tissue spherical deconvolution^110^ and subsequently underwent multi-tissue informed log-domain intensity normalisation. Anatomically constrained tractography was performed systematically by generating streamlines using second order integration over fibre orientation distributions with dynamic seeding^111,112^. Streamline generation was aborted when 40 million streamlines had been accepted. Using a spherical-deconvolution informed filtering of tractograms (SIFT2) approach, interregional tract strength (TS) was taken as the streamline count weighted by the estimated cross section^112^. A group-representative 200×200 TS matrix was generated using distance dependent consensus thresholding^113^. The approach involves varying the consensus threshold as a function of distance. The resulting connectivity matrix preserves the pooled edge length distribution of subject-level data as well as integral organisational features, such as long-range connections, while reducing false positive edges. The group-representative matrix contained 13.7% of possible edges.

##### Correspondence of cortical wiring features

To assess the complementariness of these features in characterising cortical wiring, we computed matrix-wide Spearman correlations between all feature pairs (MPC-GD, MPC-TS, GD-TS). We also assessed regional variations in feature correspondence at each node using Spearman correlations and the standard deviation in a multi-feature fingerprint (**Figure S1**).

#### d) Building the wiring space

##### Overview of approach

The wiring space was built through the integration of MPC, GD, and TS. In an effort to provide our community access to the methods we used here, we have made normative manifold maps openly available (https://github.com/MICA-MNI/micaopen/tree/master/structural_manifold) and incorporated all relevant functions and workflow into the BrainSpace toolbox (http://brainspace.readthedocs.io^114^). The procedure is as follows:

i. **Normalisation:** Nonzero entries of the input matrices were rank normalised. Notably, rank normalisation was performed on the inverted form of the GD matrix *i.e.,* larger values between closer regions. The less sparse matrices (GD and MPC) were rescaled to the same numerical range as the sparsest matrix (TS) to balance the contribution of each input measure.
ii. **Fusion:** Horizontal concatenation of matrices and production of a node-to-node affinity matrix using row-wise normalised angle similarity. The affinity matrix thus quantifies the strength of cortical wiring between two regions. Alternative data fusion techniques, such similarity network fusion^115^ and joint embedding^64^, aim to identify similar motifs across modalities. A key outcome of those approaches is higher signal-to-noise ratio, however, unique network information provided by each modality would be minimised. Given our cross-modal structural analyses highlighted special principles of cortical organisation are represented by each modality, we sought to use the concatenation approach that preserves distinct information in each modality.
iii. **Manifold learning:** Diffusion map embedding was employed to gain a low dimensional representation of cortical wiring. Diffusion map embedding belongs to the family of graph Laplacians, which involve constructing a reversible Markov chain on an affinity matrix. Compared to other nonlinear manifold learning techniques, the algorithm is robust to noise and computationally inexpensive^116,117^. A single parameter α controls the influence of the sampling density on the manifold (α = 0, maximal influence; α = 1, no influence). As in previous studies^36,118^, we set α = 0.5, a choice retaining the global relations between data points in the embedded space. Notably, different alpha parameters had little to no impact on the first two eigenvectors (spatial correlation of eigenvectors, r>0.99). We operationalised a random walker to approximate the likelihood of transitions between nodes, illuminating the local geometry in the matrix. Preservation of local geometry using the kernel critically differentiates diffusion maps from global methods, such as principle component analysis and multidimensional scaling. Local geometries are integrated into a set of global eigenvectors by running the Markov chain forward in time. The decay of an eigenvector provides an integrative measure of the connectivity between nodes along a certain axis. This lower dimensional representation of cortical wiring is especially interesting for interrogating the cortical hierarchy, which previous research suggests extends upon sensory-fugal and anterior-posterior axes. In the present study, the number of dimensions selected for further analysis was determined based on the variance explained by each eigenvector, where the cut-off point determined using the Cattell scree test. This resulted in two dimensions, which aligns with the hypothesised number of axes and, fortunately, can be readily interpreted.

##### Key outcome metrics

The wiring space represents the principle axes of variation in cortical wiring, as well as their interaction. We displayed the conversion from anatomical to wiring space using a three-part colourmap. The colour of each node was ascribed based on proximity to the limits of the wiring space; blue for closeness to the maximum of the first eigenvector, green for closeness to minimum of the first eigenvector, and redness represents closeness to maximum of the second eigenvector.

The relative positioning of nodes in the wiring space informs on the strength of cortical wiring along the principle axes. We characterised the relative positioning of each pair of nodes with wiring distance and difference along each primary axis, which pertain to node-to-node proximity and axis-specific shifts, respectively. To calculate wiring distances, we triangulated the wiring space used a Delaunay approach and calculated geodesic distance between each node used the Fast Marching Toolbox (https://github.com/gpeyre/matlab-toolboxes/tree/master/). The average wiring distance of each node informs upon centrality within the space and reflects a region’s propensity to have many cortical connections.

### Neurobiological substrates of the wiring space

#### a) Association to cytoarchitectural features

For the cytoarchitectonic maps, a 100μm resolution volumetric histological reconstruction of a *post mortem* human brain from a 65-year-old male was obtained from the open-access BigBrain repository^8^, on February 2, 2018 (https://bigbrain.loris.ca/main.php). Using previously defined surfaces of the layer 1/11 boundary, layer 4 and white matter^119^, we divided the cortical mantle into supragranular (layer 1/11 to layer 4) and infragranular bands (layer 4 to white matter). Staining intensity was sampled along five equivolumetric surfaces within the predefined supra- and infra-granular bands at 163,842 matched vertices per hemisphere, then averaged for each parcel. We estimated cytoarchitectural similarity of regions by performing the above MPC procedure on BigBrain derived intracortical profiles, as in previous work^7^. Externopyramidisation^45^, described as the “gradual shift of the weight of the pyramidal layers from the V to the IIIc,” was approximated as the product of the normalised peak intensity and the relative thickness of the supragranular layers, *i.e.*

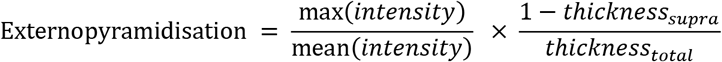

Intensity and thickness quotients were independently rescaled between 0 and 1 across all regions to balance their contribution to the externopyramidisation metric. Higher values reflect higher intensity values and shallower depth of the peak layer.

#### b) Cell-type specific gene expression

Cell-type specific gene lists were derived from an analysis of >60,000 single cells extracted from human adult visual cortex, frontal cortex, and cerebellum with single-nucleus Droplet-based sequencing (snDrop-seq) or single-cell transposome hypersensitive-site sequencing (scTHS-seq)^120^. We focused on eight canonical cell classes; astrocytes, endothelial cells, microglia, inhibitory neurons, excitatory neurons, oligodendrocytes, OPCs, and pericytes. Cell-type specific expression maps were calculated as the average of log2 normalised gene expression across eleven neocortical areas in twelve human adult brains^9,44^. Areas were visually matched to the nearest parcel. Inter-regional co-expression was calculated as the inverse of Euclidean distance between cell-type specific gene expression.

The influence of neurobiological similarities on relative positioning of nodes in the wiring space was tested by performing Spearman correlations of wiring distance with cytoarchitectural similarity and cell-type specific gene coexpression patterns, with and without controlling for geodesic distance. We used multiple linear regressions to evaluate the variance explained in externopyramidisation and cell-type specific expression by the two wiring space eigenvectors. Significance values were corrected to account for spatial autocorrelation in the eigenvectors using spin testing and Moran spectral randomisation, respectively^121,122^, and was operationalised using BrainSpace^114^. Spectral randomisation was initialised using the geodesic distance matrix. As an additional control analysis, we tested the correspondence of the first two eigenvectors with features of cortical morphometry using the same procedure.

The wiring space offers a dimensional approach to evaluate the concordance of gradients at multiple biological scales. To facilitate multi-scale comparisons, we generated 32 axes within the wiring space by creating inter-cardinal lines in 5.625° steps. Linear polynomial equations, corresponding to each inter-cardinal line, were evaluated for 100 equally spaced x and y-values between the minimum and maximum range of the first and second eigenvectors, respectively. Nodes were assigned the values of the nearest point along each inter-cardinal line, based on Euclidean distance, thus the position of a node on each axis could be represented by an integer (1-100). The dominant axis of variation of any feature in wiring space can be classified as the axis of maximum correlation. P-values from the Spearman correlations were subjected to FDR correction to assess whether the dominant axis of variation was significant^123^.

### Association with functional MRI based connectivity

#### Functional architecture

The wiring space was reimagined as a completely dense network with edges weighted by wiring distance (**Figure 3B**). We mapped seven established functional communities^49^ into the group-level wiring space by assigning each node to the functional community that was most often represented by the underlying vertices.

#### Predicting functional connectivity

Individual functional connectomes were generated by averaging preprocessed timeseries within nodes, correlating nodal timeseries and converting them to z scores. For each individual, the four available rs-fMRI scans were averaged at the matrix level, then the connectomes were averaged within the *Discovery* and *Hold-out* samples separately. We estimated the variance explained in functional connectivity by the cortical wiring scheme using boosted regression trees^124^. Boosted regression trees produce a predictive model from the linear weighted combination of weaker base learners that each fit the mean response of a subsection of the predictor space. Weak estimators are built in a step-wise manner, with increasing focus on poorly explained sections of the predictor space. Optimisation of the learning rate and number of estimators is critical to model complex nonlinear relationships, implicitly model interactions between predictors, and reduce overfitting. Overfitting was further reduced and predictive performance enhanced by using a random subset of data to fit each new tree. The present study specifically used the AdaBoost module of scikit-learn v0.21.3in Python3.5 and established the optimal number of estimators and the learning rate using internal five-fold cross-validation [maximum tree depth = 4, number of estimators = 6:2:20, learning rate = (0.01, 0.05, 0.1, 0.3, 1); see **Figure S5B** for node-wise hyperparameters]. We aimed to predict functional connectivity independently for each node based on wiring distance, difference along the first eigenvector and difference along the second eigenvector to all other nodes. Each feature was z-standardised before being entered in the model. Predictive accuracy was assessed as the mean squared error and R^2^ coefficient of determination of empirical and predicted functional connectivity. Given prior z-standardisation of features, a mean squared error of 1 would represent an error of one standard deviation from the true value. An R^2^ above 0 indicates predictive value of the model, where 1.0 is the maximum possible score.

The procedure was repeated using functional connectivity across different tasks to demonstrate the flexibility of the wiring space in governing varied states of functional organisation. In line with previous work^125^, minimally-preprocessed^98^ timeseries from each of the seven tasks were averaged within node, z-standardised, then concatenated together. Functional connectivity was estimated across all tasks as the inverse of Euclidean distance, because it is less biased by outliers than Pearson correlation, which are likely to occur during active task periods. Group-average task-based functional connectivity matrices were generated for the *Discovery* and *Hold-out* groups.

### Replication of the wiring space in an independent dataset

Independent replication was performed using locally acquired data from 40 healthy adults (*MICs* cohort; 14 females, mean±SD age=30.4±6.7, 2 left-handed) for whom quantitative T1 relaxation time mapping (qT1) images were available. All participants gave informed consent and the study was approved by the local research ethics board of the Montreal Neurological Institute and Hospital. MRI data was acquired on a 3T Siemens Magnetom Prisma-Fit with a 64-channel head coil. A submillimetric T1-weighted image was acquired using a 3D-MPRAGE sequence (0.8mm isotropic voxels, 320×320 matrix, 24 sagittal slices, TR=2300ms, TE=3.14ms, TI=900ms, flip angle=9°, iPAT=2) and qT1 data was acquired using a 3D-MP2RAGE sequence (0.8mm isotropic voxels, 240 sagittal slices, TR=5000ms, TE=2.9ms, TI 1=940ms, T1 2=2830ms, flip angle 1=4°, flip angle 2=5°, iPAT=3, bandwidth=270 Hz/px, echo spacing=7.2ms, partial Fourier=6/8). The combination of two inversion images in qT1 mapping minimises sensitivity to B1 inhomogeneities^42^, and provides high intra-subject and inter-subject reliability^126^. A spin-echo EPI sequence was used to obtain diffusion weighted images, consisting of three shells with b-values 300, 700, and 2000s/mm2 and 10, 40, and 90 diffusion weighting directions per shell, respectively (TR=3500ms, TE=64.40ms, 1.6mm isotropic voxels, flip angle=90°, refocusing flip angle=180°, FOV=224×224 mm^2^, slice thickness=1.6mm, mb factor=3, echo spacing=0.76ms). One 7 min rs-fMRI scan was acquired using multi-band accelerated 2D-BOLD echo-planar imaging (TR=600ms, TE=30ms, 3mm isotropic voxels, flip angle=52°, FOV=240×240mm^2^, slice thickness=3mm, mb factor=6, echo spacing=0.54mms). Participants were instructed to keep their eyes open, look at fixation cross, and not fall asleep.

The data preprocessing and construction of the wiring space were otherwise virtually identical to the original Human Connectome Project dataset, with a few exceptions. Microstructure profiles were sampled from qT1 images. Cortical surface estimation via FreeSurfer utilised two T1 scans, and surface models were manually edited for accuracy. All fMRI data underwent gradient unwarping, motion correction, fieldmap-based EPI distortion correction, brainboundary-based registration of EPI to structural T1-weighted scan, non-linear registration into MNI152 space, and grand-mean intensity normalization. The rs-fMRI data was additionally denoised using an in-house trained ICA-FIX classifier^103,127^ as well as spike regression. Timeseries were sampled on native cortical surfaces and resampled to fsaverage via surface-based registration.

### Intracranial EEG analyses in epileptic patients

A group of ten patients with drug-resistant focal epilepsy (one male, mean±SD age=28.9±7.9, all right-handed) were scanned using the same imaging protocol as the heathy controls from the *Replication* dataset. Patients furthermore underwent intracerebral stereo-electroencephalographic investigation as part of their presurgical evaluation, after the imaging. The protocol received prior approval from the MNI Institutional Review Board. The recordings were acquired with Nihon Khoden EEG amplifiers at a sampling rate of 2000Hz, using 1 single type of depth electrodes (DIXI electrodes with either 10 or 15 electrode). A board certified neurophysiologist (BF) selected epochs without ictal events, absent of artefacts, from periods of resting wakefulness with eyes closed during standardised conditions, resulting in 1-2 minutes of recording for each patient.

Each depth electrode was mapped to a cortical parcel using with the following procedure. For each subject, cortical surfaces were extracted from the high-resolution pre-implantation T1-weighted using FreeSurfer6.0. Next, a clinical structural image, acquired during the implantation period on a Philips Medical Systems 1.5T MRI scanner, was transformed to the T1-weighted space using volume-based affine transformation with nearest neighbour interpolation. For seven patients, the clinical scan was a T1-weighted image (T1_3D_SENSE, slice thickness=0.78mm, number of slices=280, single echo, phase encoding steps=320, echo train length=320, TR=0.0079s, flip angle=6°, multi coil receiver coil, TE=0.0035s). For three subjects, the clinical scan was an Axial T2 scan (slice thickness=2mm, number of slices=242, single echo, echo train length=141, TR=2.8, flip angle=90°, multi coil receiver coil, TE=0.48s). Using tissue-type specific maps and individualised surface reconstructions, each electrode in grey matter was mapped to the nearest surface vertex, and labelled as the corresponding parcel, based on minimum geodesic distance from the centroid coordinate of the electrode to the cortical midsurface.

### Directed information processing

Intracranial EEG signals were re-referenced to the average signal of white matter channels to remove scalp reference and suppress far-field potentials caused predominantly by volume conduction^128^. The auto spectral density of each channel, *P_xx_*, and the cross power spectral density between pairs of within-subject channels, *P_xy_*, were calculated with Welch’s method (59 overlapping blocks, 2s duration, 1s steps, weighted by Hamming window)^129^. These measures allow for the calculation of magnitude squared coherence between two signals^130,131^

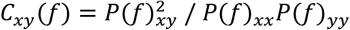

In the above formula, P_xx_ and P_yy_ are power spectral density estimates and Pxy is the cross spectral density estimate. Coherence was evaluated in 0.77 Hz steps (n=129) from 0.5-100Hz. The 55-65 Hz range was not inspected due to power line noise at 60 Hz. We used boosting regression models to estimate variance explained in undirected coherence by the wiring space [see *Predicting Functional Connectivity* for details]. In contrast to the fMRI-analysis, however, only within-sample variance explained (R^2^) was examined.

The temporal coupling of two signals was determined by the phase-slope index, using the formula:

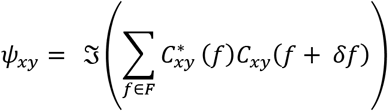

In the above formula, C_xy_ is complex coherence as defined above, δf is the frequency resolution, 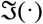 denotes taking the imaginary part and F is the set of frequencies over which the slope is summed^51^. We used a sliding window approach for defining frequency bands, using 4Hz bandwidth and 2Hz overlap from 1-100Hz, excluding the 55-65Hz range due to power line artefact. The phase slope index leverages the relationship between increasing phase difference with increasing frequency to establish the driver and respondent sources^51^. Given incomplete coverage of intracranial electrodes in each subject, we discretised the wiring space into a set of sub-sections using consensus k-means clustering. Consensus-based k-means clustering and converged on a stable solution of k=12 across 100 repetitions, provided the k range of 10-20^132^. 11 out of 12 clusters were represented in the intracranial data (**Figure S7**). We used a linear mixed effect models to approximate the relationship between phase-slope index within each frequency band and cluster membership:

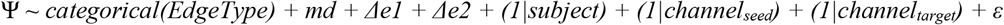

Here, Ψ stands for phase slope index and *EdgeType* was defined by the seed and target clusters, in such that the *EdgeType* of a connection from cluster 1 to cluster 2 would be ‘12’. *EdgeType* was a fixed effect, *subject* and *channels* were nested in the model with random intercepts. We also included wiring distance (*md*), difference on eigenvector 1 *(Δe1)* and difference on eigenvector 2 *(Δe2)* as fixed effects to account for variations in the positioning on nodes within a cluster. We used the t-statistics of each *EdgeType* category as a measure of phase slope index between clusters, then vectorised the t-statistics across all frequency bands for each cluster and used Pearson correlations to estimate the similarity of cluster’s directed coherence patterns. For each cluster-to-cluster correlation, the phase slope index of that direct relationship was removed, thus the correlation indicates the similarity of phase slope index to all other clusters. A principle component analysis was used to extract the main axes of variation in the cluster similarity matrix. This component loading was cross-referenced with cell-type specific gene expression, with regions labelled by the corresponding cluster, as well as average externopyramidisation estimates for each cluster. Due to the limited number of observations to predictor variables, we opted for lasso regularisation and focused on high regularisation/sparsity models to characterise features importance^133^. The standard deviation in fitted least-squares regression coefficients was calculated using a leave-one-observation-out procedure, all of which used the range of lambdas from the full model. We performed a post-hoc multiple linear regression with the most sparse model to evaluate the variance explained in the component loading by few cellular features (adjusted R^2^). Next, we identified the most influential frequency bands to the component loading by performing Pearson correlations between the average phase slope index spectra with the component loading scores. Inspecting the frequency bands with maximum and minimum rho values, we performed significance thresholding of the cluster-to-cluster t-statistic matrices using the fixed effect p-values from the linear mixed effect model with an alpha level of 0.05. Standard deviations in the t-statistics were quantified across leave-one-subject-out iterations to ensure robustness of the direction and strength of the phase slope index estimates. Finally, we followed the criteria set forth by Felleman and van Essen (1991) to test whether the frequency-specific phase slope index networks conformed to a hierarchical topology. We performed this in a “top-down” fashion by progressively adding clusters to lower levels of the model based on the driver-respondent relationships shown in the thresholded t-statistic matrices. First, clusters that only drive oscillations *(i.e.,* positive t-statistics) were placed at the top level of the hierarchy, then the next level was populated by clusters that only respond to clusters in upper levels, and so forth. The internal consistency of the hierarchy is determined by whether all significant edges can be placed into the model with a constant flow of directed coherence from top to bottom. Preprocessing was performed using the FieldTrip toolbox^134^, while the cross-spectral density estimates, phase slope index (http://doc.ml.tu-berlin.de/causality/) and linear models were estimated using MATLAB^135^.

### Code and data availability statements

Preprocessed group-level matrices, normative manifold maps and integral scripts are openly available at https://github.com/MICA-MNI/micaopen/tree/master/structural_manifold

## Acknowledgements

The authors would also like to express their gratitude to the open science initiatives that made this work possible, including the teams involved in the BigBrain project, the PsychEncode consortium, the Human Connectome Project and Scikit-learn. Furthermore, we thank Dr. Matthias Kirschner for his helpful insight on the manuscript, and the MRI and EEG technicians at the Montreal Neurological Institute.

## Notes

https://github.com/MICA-MNI/micaopen/tree/master/structural_manifold

### Competing Interest Statement

The authors have declared no competing interest.

