## Supplementary material for "A cortical wiring space links cellular architecture, functional dynamics and hierarchies in humans": Figure S

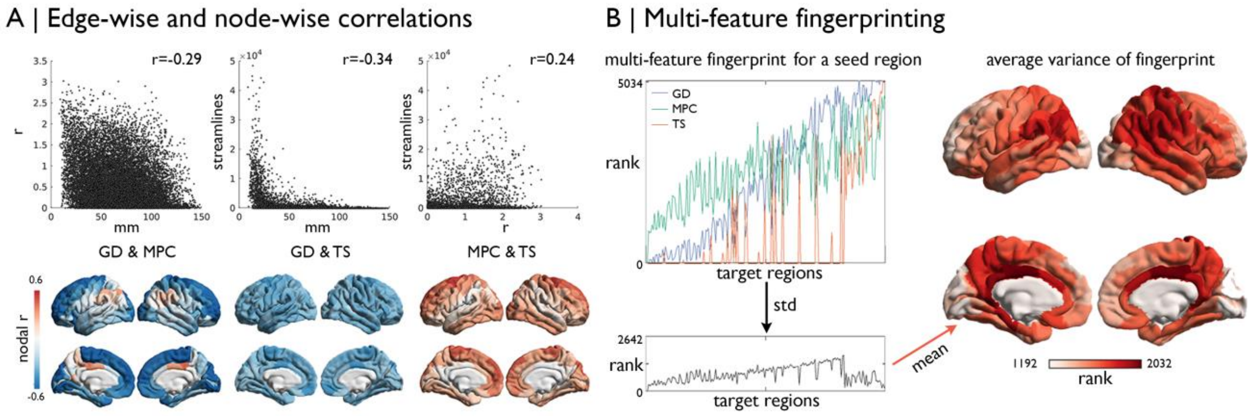


**Figure S1: (A)** Scatterplots depict the global correlation between the different cortical wiring features *i.e.*, geodesic distance (GD), microstructure profile covariance (MPC), and diffusion-based tract strength (TS). Average r values for each node are projected below on the cortical surface. **(B)** Fingerprinting involved combining all three wiring features of one region. The standard deviation across the three features was estimated for each edge, then the average was taken for each node as a measure of feature variance of the fingerprint.

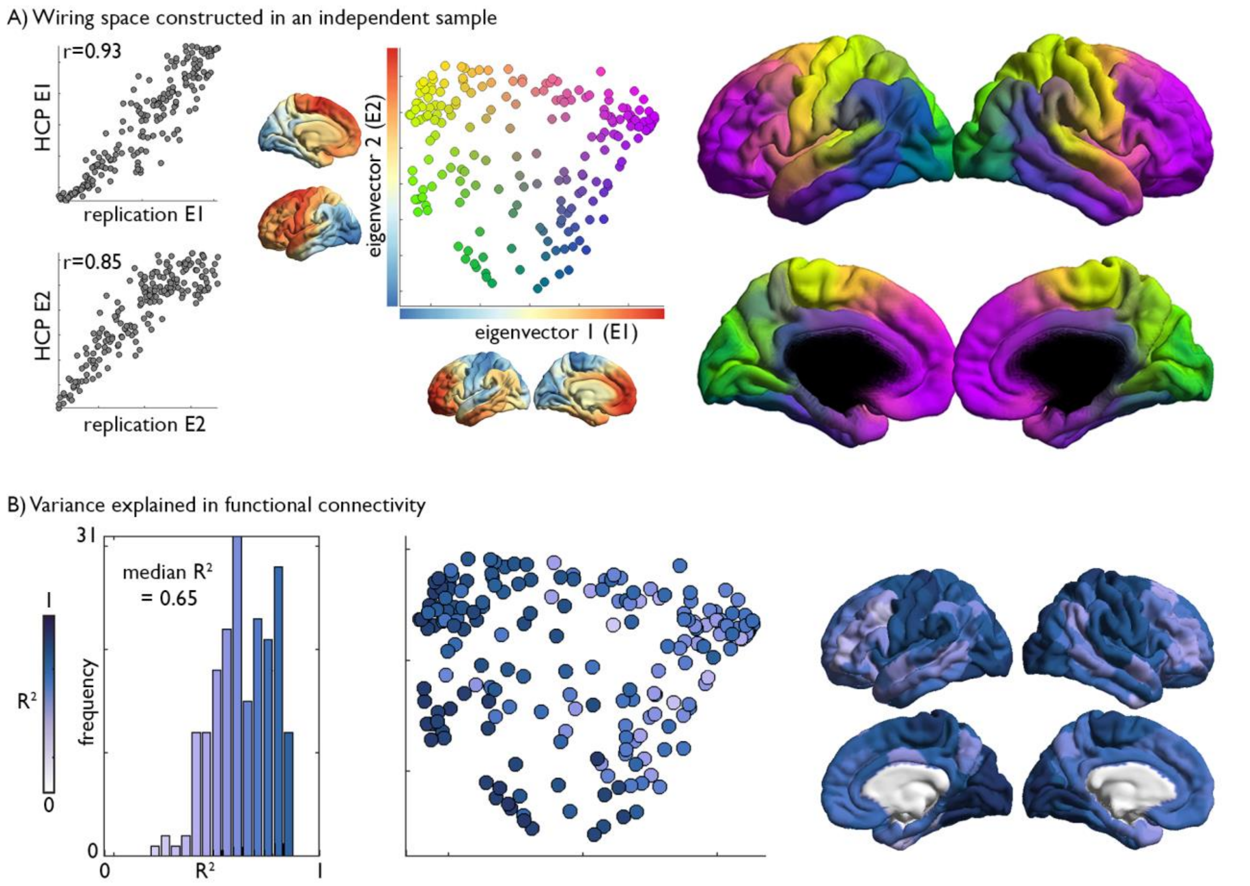


**Figure S2: Independent replication of the wiring space.** The wiring space was regenerated using an independent *MICs* dataset of 40 healthy adults scanned at our imaging centre. Imaging parameters were similar to the original cohort, albeit using quantitative T1 relaxometry as a marker of cortical microstructure rather than HCP's T1w/T2w ratio mapping. **(A)** Scatterplots show marked correspondence between the original HCP *Discovery* sample eigenvectors and those from the *MICs* dataset, with Spearman correlations shown. **(B)** Using boosting regression, we similarly found that high, but regionally variable, variance in the group-average functional connectivity could be explained by the wiring space in the *MICs* dataset.

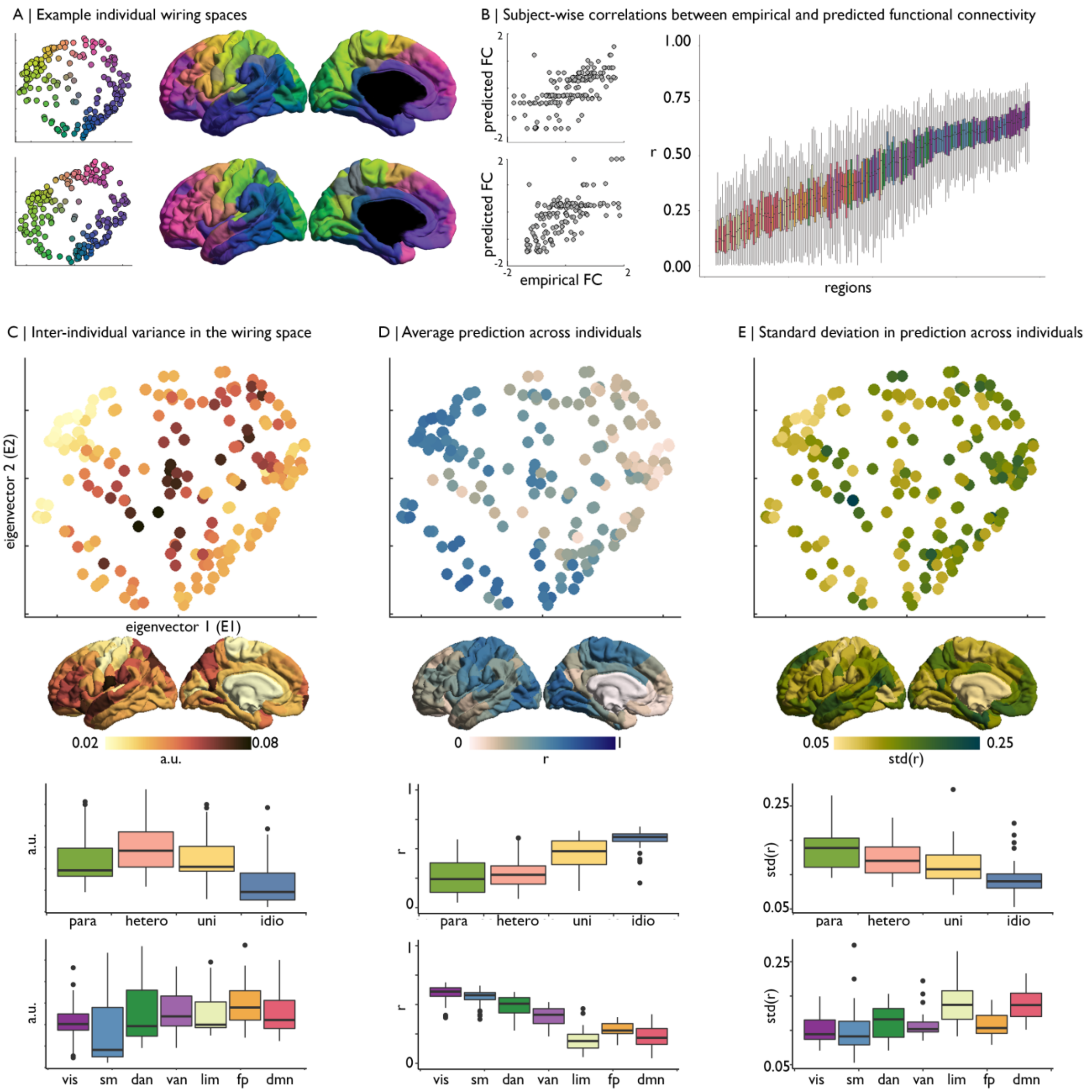


**Figure S3: Individual variation in the wiring space and prediction of rs-fMRI connectivity. (A)** Wiring spaces constructed in two individuals of the *Hold-out*, which were aligned to the group-level *Discovery* wiring space. **(B)** Correlation between predicted and empirical functional connectivity. Each box represent a node and the distribution is taken across individuals. Box are coloured by the assignment of the node to a functional community (Figure 3). **(C-E)** Individual variation was calculated as the Euclidean distance between the node's position in the subject vs group-average wiring space, where the position is synonymous with the values on the first two eigenvectors following Procrustes alignment. Inter-individual variance was taken as the average across all subjects and provided as arbitrary units (a.u.). We also calculated the mean and SD in the correlation between predicted and empirical functional connectivity for each node across subjects. Node-wise values are presented in the wiring space, on the cortical surface and stratified by level of laminar differentiation[7,136] and functional network[49].

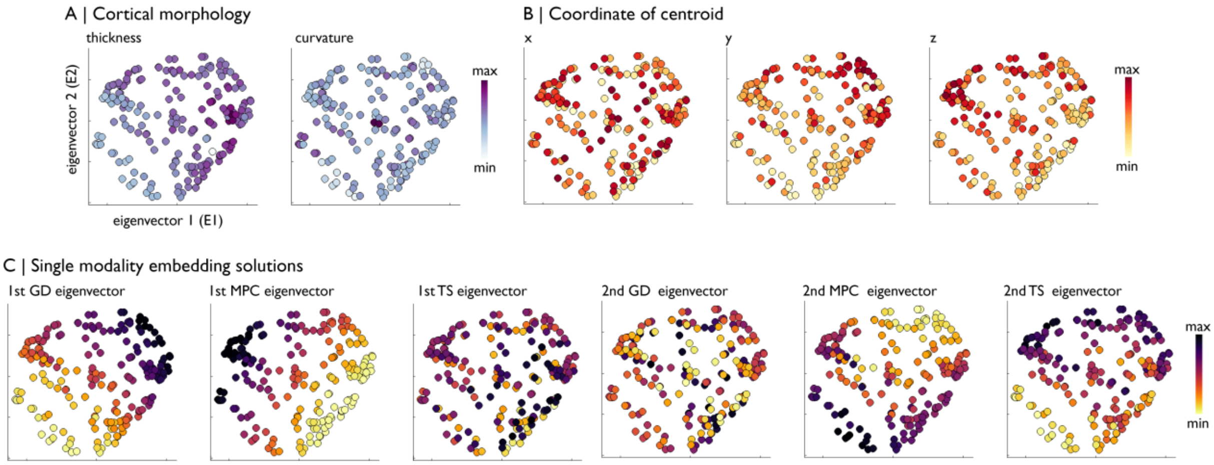


**Figure S4: Wiring space captures the interplay of more simple axes. (A)** Group-average cortical thickness and curvature measures in the manifold. **(B)** Centroid coordinate of each parcel. x=left-right, y=posterior-anterior, z=inferior-superior. **(C-D)** Principles gradients from single modalities.

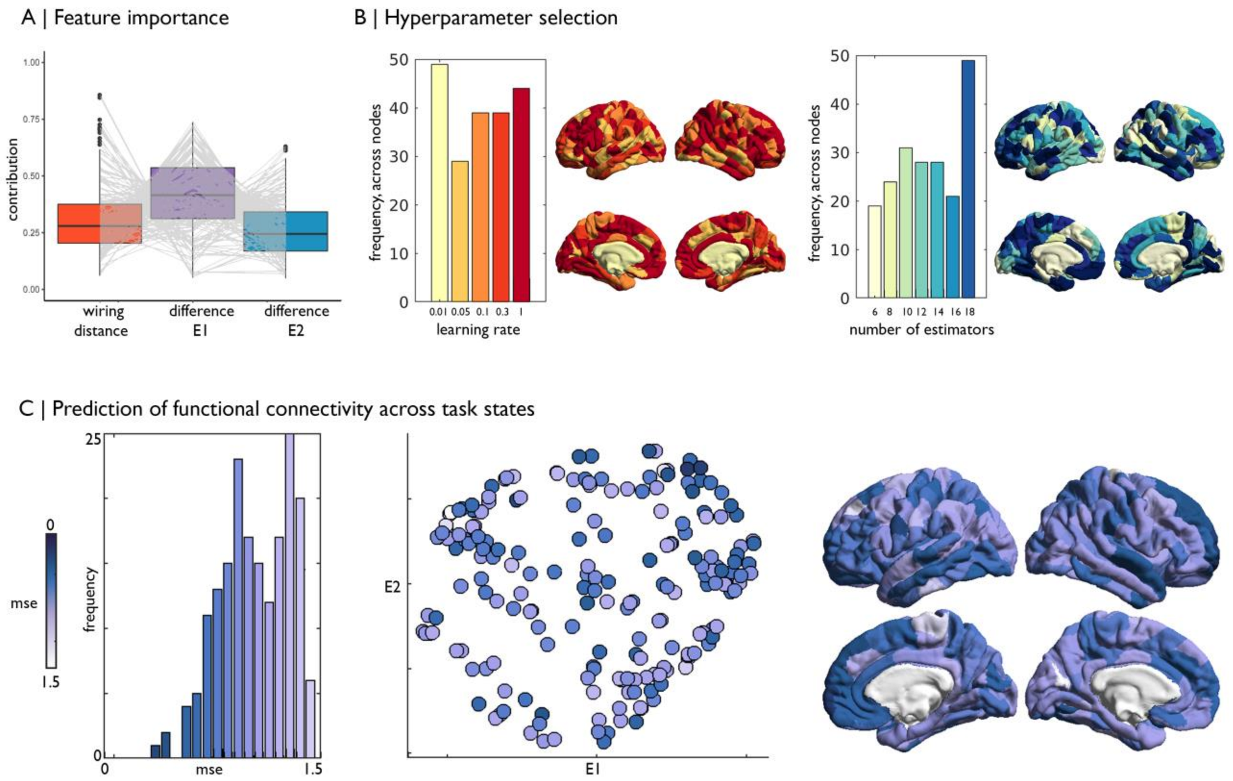


**Figure S5: Modelling fMRI-derived connectivity from the wiring space. (A)** Box and spaghetti plots depict the relative importance of wiring space features for the boosting regression model at each node. **(B)** Learning rate and number of estimated selected by cross-validation for the model at each node. **(C)** Predictive accuracy, measured by the mean squared error, for task-related functional connectivity.

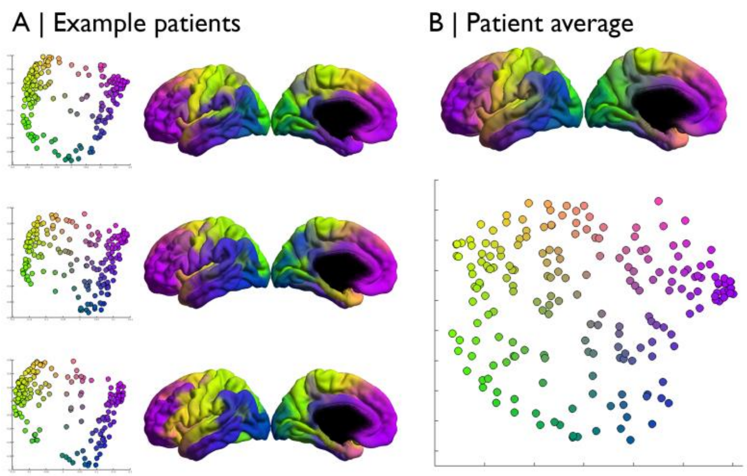


**Figure S6: Patient wiring space**. **(A)** Wiring spaces were generated for each of the ten patients. **(B)** The wiring space generated from the group-average of cortical wiring features across the patients was highly similar to the healthy template (correlations between eigenvectors 1/2: r=0.83/0.82).

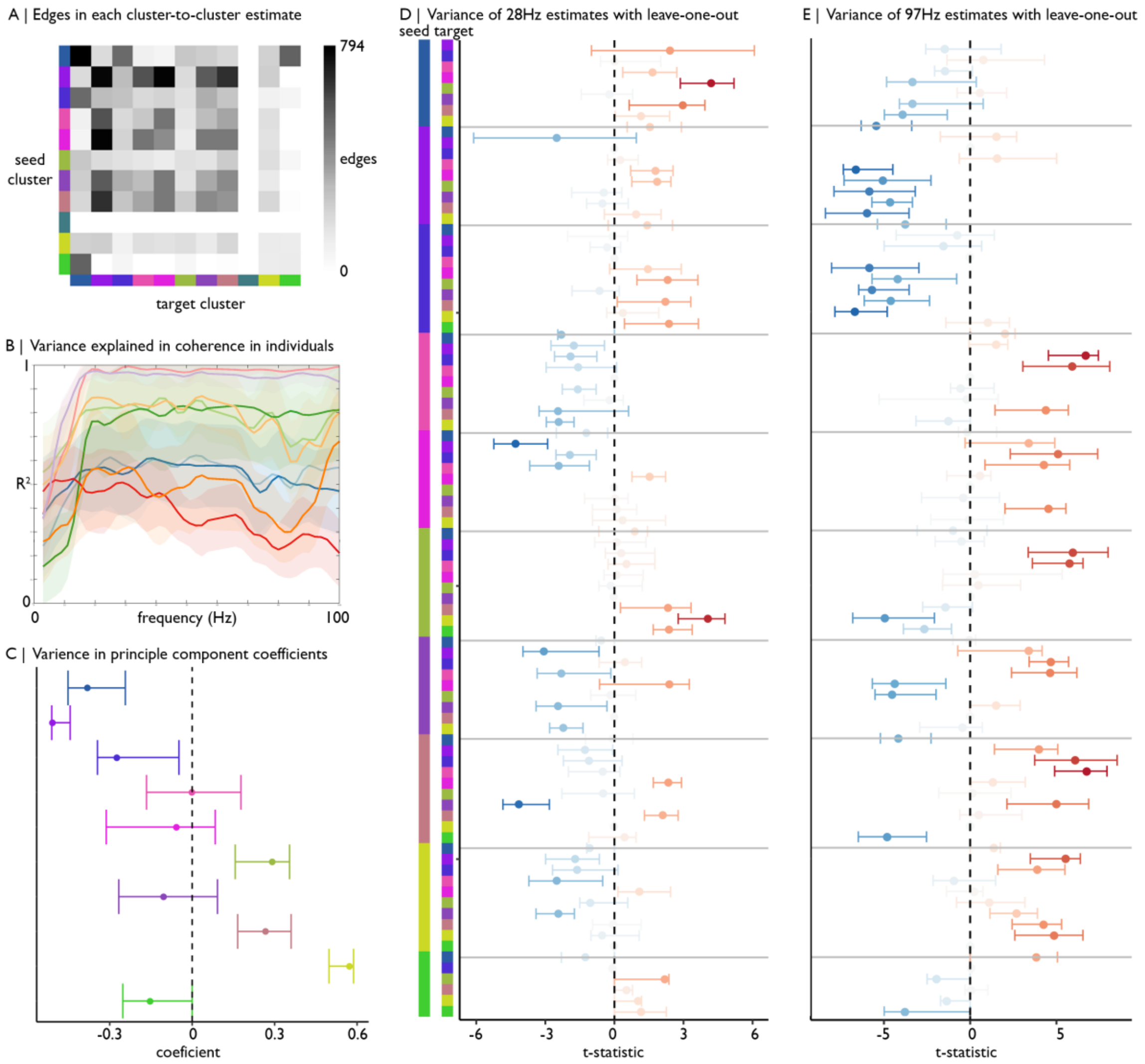


**Figure S7: Robustness of intracranial EEG analyses. (A)** Matrix depicting the number of intra-subject electrode pairs that contribute to the cluster-to-cluster estimations of phase slope index. **(B)** Variance explained in coherence by boosting regression, operationalised at a single subject level with individualised wiring spaces. **(C)** Variance in coefficients from a principle component analysis of inter-cluster similarities in the phase slope index using a leave-one-subject-out procedure. **(D)** and **(E)** involve the same leave-one-subject-out procedure to gauge the variance in edge-wise phase slope index estimates at frequencies of interest. Notably, the range of estimates does not pass zero for significant edges.

**Table S1**: Variance explained in externopyramidisation with alternative input measures

| | Adjusted $R^2$ |
|---|---|
| GD only | 0.50975 |
| MPC only | 0.49381 |
| TS only | 0.62106 |
| GD + MPC | 0.64716 |
| GD + TS | 0.52581 |
| MPC + TS | 0.54945 |
| GD, MPC + TS | **0.65076** |

**Table S2: Statistical relationships between cell-type specific gene expression and wiring space.**

| | Regional expression modelled by wiring space eigenvectors 1 & 2[1] | | Spearman correlation between co-expression and wiring distance | | | |
|---|---|---|---|---|---|---|
| | | | Full correlation | | Corrected for geodesic distance | |
| Cell-type | r | p[2] | r | p | r | p |
| Astrocyte | 0.91 | 0.0019 | -0.74 | <0.001 | -0.48 | 0.0007 |
| Endothelial cells | 0.81 | 0.2817 | -0.55 | <0.001 | -0.28 | 0.0017 |
| Microglia | 0.76 | 0.0368 | -0.75 | <0.001 | -0.53 | <0.001 |
| Neurons – excitatory | 0 | 0.1271 | -0.68 | <0.001 | -0.42 | <0.001 |
| Neurons – inhibitory | 0.46 | 0.0404 | -0.68 | <0.001 | -0.41 | 0.0001 |
| OPCs | 0.89 | 0.0273 | -0.65 | <0.001 | -0.34 | 0.0002 |
| Oligodendrocytes | 0 | 0.3601 | -0.52 | <0.001 | -0.33 | 0.0002 |
| Pericytes | 0.35 | 0.2534 | -0.64 | <0.001 | -0.33 | 0.0002 |

[1] Expression ~ $1 + \beta_1$eigenvector1 + $\beta_2$eigenvector2+ $\varepsilon$

[2] Comparison to null model with 10000 permutations of spatial maps with matched spatial autocorrelation [121]
